# datadrivenhypothesis.org: A resource for metabolic gene discovery through integrated pathway co-essentiality mapping

**DOI:** 10.64898/2026.02.11.705287

**Authors:** Matthew D. Hirschey, Pol Castellano-Escuder, John Bradley

## Abstract

datadrivenhypothesis.org (DDH) integrates gene dependency, expression, literature data, and beyond from ∼20,000 human genes to reveal hidden metabolic connections. Through pathway-level co-essentiality analysis with novelty scoring, DDH uncovered unexpected links including Complex II coupled to purine nucleotide biosynthesis. This broad platform offers intuitive tools for hypothesis generation via gene queries, pathway analysis, and network visualization, plus publication-ready downloads. Automated updates make DDH a dynamic resource, bridging genes with unknown function to actionable metabolic insights.

## MAIN

Despite decades of research, approximately 20-50% of predicted human genes lack functional annotation ^1-3^. Most human genes have fewer than 10 publications, while a small fraction dominate the literature (Fig. 1A). Further, when we examine metabolically defined gene subsets, this pattern persists: genes associated with inborn errors of metabolism (n=1,766; IEMbase^4^) and mitochondrial genes (n=1,136; MitoCarta3.0^5^) show similar distributions (Fig. 1B-C). Binning genes by publication count reveals that across all categories, the majority of genes have few publications, highlighting a substantial opportunity for functional discovery even among metabolic disease-associated genes (Fig. 1D). Traditional gene function discovery approaches study individual genes or simple pairwise interactions. These methods fail to capture the emergent properties of metabolic pathways working in concert. The Cancer Dependency Map (DepMap) has generated CRISPR essentiality data across hundreds of cancer cell lines. This resource creates unprecedented opportunities for functional discovery. Co-essentiality mapping identifies genes sharing similar patterns of essentiality across diverse cellular contexts ^6^. This approach has successfully revealed important functional relationships. However, nearly 200 million possible gene-gene comparisons present substantial interpretation challenges. Further, focusing on individual gene queries lacks pathway-level analyses. The metabolic research community needs accessible tools that organize this complexity into biologically meaningful insights while maintaining analytical rigor.

**Figure 1:**
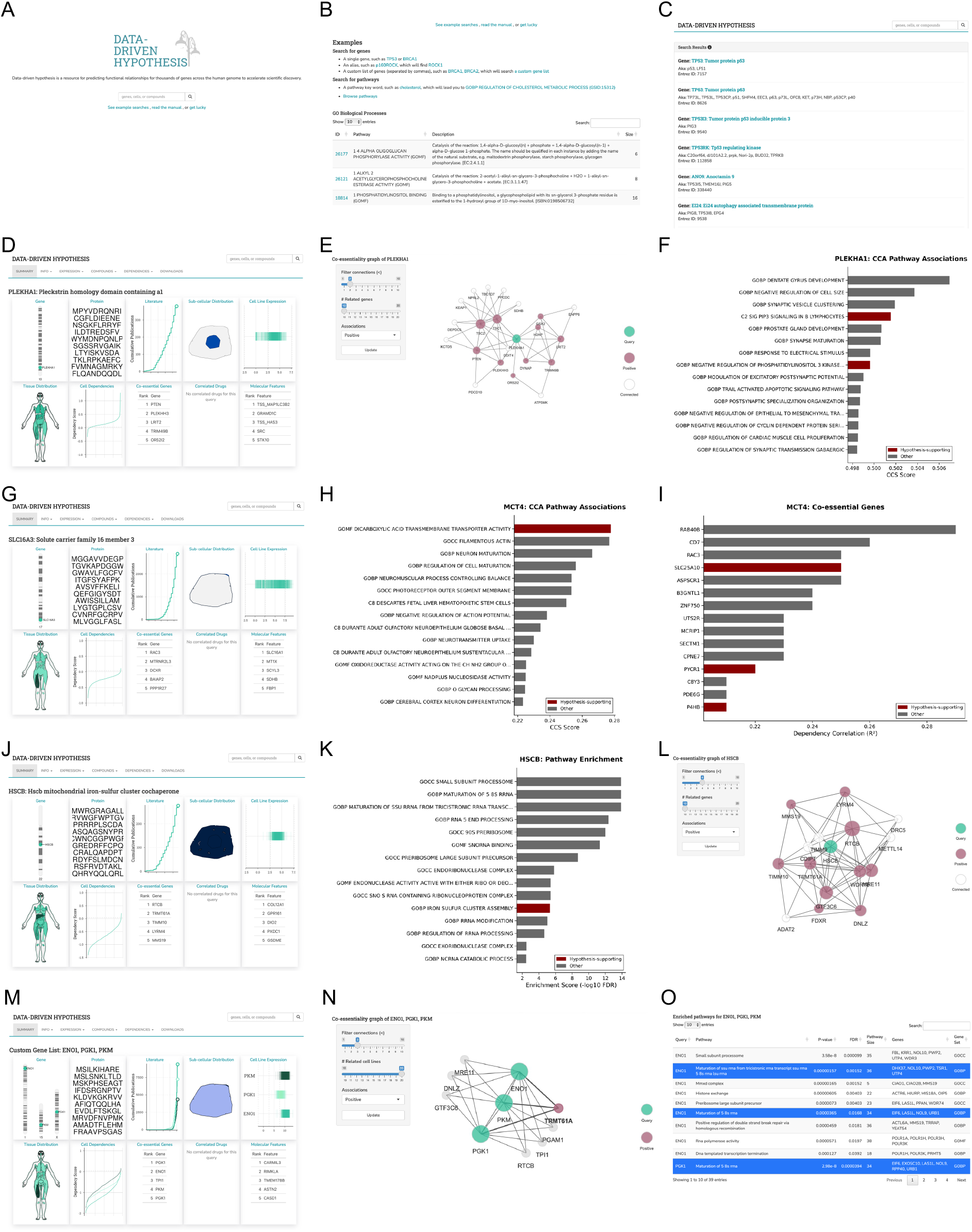
Publication frequency across human genes. **(A)** Publication counts per human gene, ranked by frequency. Most genes have few publications, while a small subset dominates the literature. **(B**,**C)** Publication counts for genes associated with inborn errors of metabolism (n=1,766; IEMbase) or mitochondrial genes (n=1,136; MitoCarta3.0). Genes with the highest number of publications are labeled. **(D)** Distribution of genes by publication bins across all three datasets. The majority of genes have few publications regardless of functional category, though disease-associated and mitochondrial gene subsets show enrichment in higher publication bins.

### Integrating multiple data dimensions for enhanced discovery

The datadrivenhypothesis.org platform addresses these challenges through a multi-layered approach to gene discovery (Figure 2A). At its core, DDH integrates several complementary core data types, including: (1) CRISPR-based gene dependency scores from 739 cell lines spanning diverse lineages, (2) matched gene expression data from 733 cell lines to filter noise from unexpressed genes, and (3) literature co-occurrence patterns from 30 million PubMed abstracts to distinguish novel from established relationships. Beyond these core data types, DDH also integrates additional NCBI/PubMed data, Protein & Structure data, Human Metabolome Database (HMDB) data, Cell Line Resource data, and other gene metadata.

**Figure 2:**
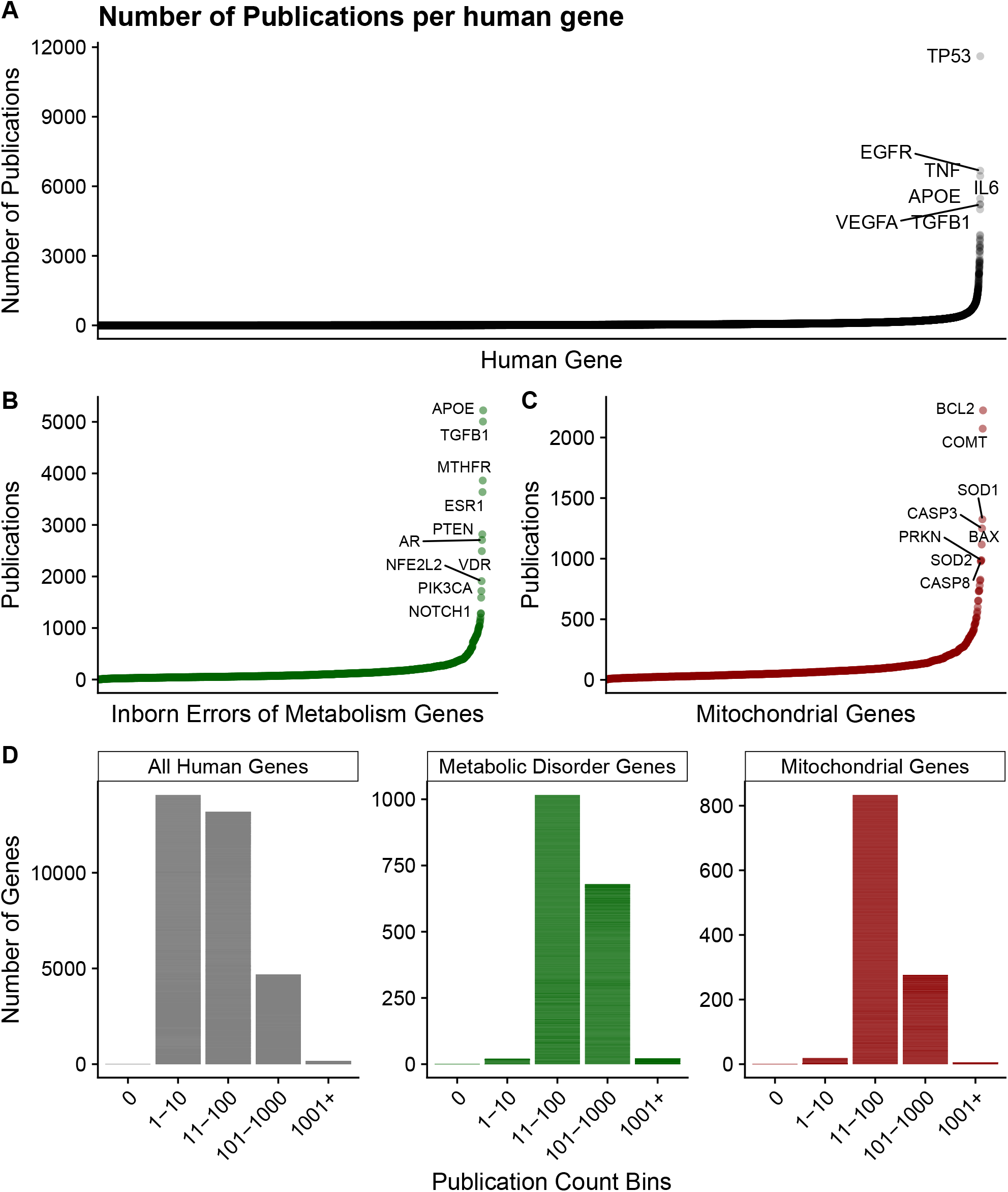
DDH platform interface and discovery examples. **(A)** Homepage with search interface for gene queries. **(B)** Example queries for single genes, pathways, and custom gene lists. **(C)** Search results display with integrated data sources. **(D-F)** PLEKHA1 query showing dependency distribution, top co-essential genes including mTOR regulators (TSC1/TSC2, PTEN), and CCA pathway associations. **(G-I)** MCT4/SLC16A3 query revealing co-essentiality with acetoin metabolism (DCXR) and glycolytic enzymes, with pathway enrichment analysis. **(J-L)** HSCB query demonstrating co-dependencies with known Fe-S cluster proteins (FDXR, FXN) and unexpected candidates including ADAT2, visualized as interaction network. **(M-O)** Custom glycolysis gene query (ENO1, TPI1, PGK1, PKM) showing coordinated dependencies, network topology with tRNA modification enzymes, and ranked pathway associations.

The user interface resembles a familiar and intuitive internet-search site (Fig. 2A), where single or multiple genes can be queried. Alternatively, example queries can be selected (Fig. 2B). Data is then presented to the user in a ‘search results’ list (Fig. 2C), which then when selected triggers data fetching and aggregation for presentation via a simple dashboard.

One key innovation lies in pathway-level co-essentiality mapping. Rather than analyzing ∼200 million individual gene pairs, we first extract Principal Components from ∼3,000 annotated pathways (MSigDB collections ^7^, including GO, KEGG, Reactome), then apply Canonical Correlation Analysis on these signatures to identify pathway-pathway relationships. This dimensionality reduction approach revealed that metabolic pathways exhibit particularly strong co-essentiality patterns, likely reflecting coordinated enzyme dependencies or shared rate-limiting steps.

Additionally, our co-publication index provides crucial context for distinguishing novel from established gene relationships. We leveraged pre-defined machine learning annotations, which assign standardized gene identifiers to all 30 million PubMed abstracts using natural language processing ^8^. For each gene pair, we count co-occurrences within individual abstracts across the entire corpus. Raw co-occurrence counts range from 0 (never published together) to 23,197 (for highly studied gene pairs like TP53-MDM2). To facilitate interpretation, we normalized these counts to a 0-100 scale, where 100 represents the maximum observed co-occurrence. This index enables researchers to immediately identify unexpected connections

### Proof-of-Concept: Discovering the Complex II-purine synthesis axis

To demonstrate DDH’s utility for metabolic discovery, we analyzed electron transport chain (ETC) complexes as an exemplar system ^9^. Hierarchical clustering of pathway co-essentiality patterns revealed an unexpected finding. Complex II (succinate dehydrogenase) clustered separately from Complexes I, III, IV, and V. These four complexes grouped together, consistent with their ability to form respiratory super-complexes. Complex II showed unique co-essentiality with three metabolic pathways absent from other ETC complexes: “Nucleobase Metabolic Process” (r = 0.72), “Purine Nucleobase Monophosphate Metabolic Process” (r = 0.68), and “Glutamine Metabolic Process” (r = 0.61). This computational prediction led to extensive experimental validation in our recent study ^9^. Complex II inhibition in AML cells reduced de novo purine biosynthesis by 60%. Exogenous purines (adenine/guanine) fully rescued viability following Complex II blockade. The mechanism involves glutamate accumulation, disrupting nitrogen donation for purine ring assembly. High SDHB expression correlates with venetoclax resistance in AML patients (Beat AML dataset). This discovery established Complex II as a metabolic vulnerability in AML with immediate therapeutic implications. DDH’s pathway-level analysis enabled this finding.

### Future Discovery Opportunities

The DDH platform enables data-driven discovery by revealing unexpected metabolic connections hidden within functional genomics data. Our validation of the Complex II-purine metabolism link described above provides a roadmap for how computational predictions from DDH can lead to transformative biological insights. Here, we highlight additional metabolic hypotheses emerging from pathway-level analysis that await experimental validation.

Several intriguing metabolic connections emerge from DDH analysis:

- **mTOR signaling regulation through phosphoinositide metabolism**: PLEKHA1 is a pleckstrin homology domain protein that binds PI(3,4)P2 (Fig. 2D). This gene shows strong co-essentiality with TSC1/TSC2 complex members and PTEN (Fig. 2E). Known mTOR regulators, including DEPDC5, NPRL2, and the TSC complex components, all appear in the top co-essential genes (Fig. 2E). The hypothesis: PLEKHA1 acts as a PI(3,4)P2 sensor that inhibits mTOR by sequestering this lipid, perhaps from AKT activation sites (Fig. 2F). Testing this would require measuring mTOR activity (pS6K levels) after PLEKHA1 knockdown under normal and starvation conditions.
- **Acetoin metabolism and transport**: MCT4 (SLC16A3) is a proton-linked monocarboxylate transporter that facilitates the export of lactate and pyruvate across plasma membrane (Fig. 2G, H). This gene demonstrates strong co-essentiality with diacetyl reductase (DCXR), which converts diacetyl to acetoin, and pyruvate metabolism enzymes, including PYCR1 (Fig. 2I). Multiple known glycolytic genes appear as positive controls in this network. The hypothesis: MCT4 transports acetoin, a metabolite and electron acceptor produced during fermentation. A transport assay using radiolabeled acetoin in MCT4-expressing versus control cells would directly test this prediction.
- **Iron-sulfur cluster protein identification**: HSCB is a mitochondrial DnaJ-type co-chaperone that facilitates iron-sulfur cluster biogenesis by stimulating the ATPase activity of mitochondrial Hsp70 (mortalin/HSPA9), essential for the function of Fe-S enzymes like succinate dehydrogenase and aconitase (Fig. 2J). Analysis of HSCB reveals co-dependencies with known Fe-S proteins ferredoxin reductase (FDXR) and frataxin (FXN) as positive controls (Fig. 2K), alongside unexpected hits like adenosine deaminase (ADAT2) (Fig. 2L). The hypothesis: ADAT2 and other co-essential proteins contain unrecognized iron-sulfur clusters. EPR spectroscopy or iron-55 labeling of immunoprecipitated ADAT2 would reveal Fe-S cluster presence.
- **Glycolysis-tRNA modification coupling**: Metabolic pathways, like glycolytic enzymes, can be queried together (Fig. 2M). Known glycolytic enzymes (PGK1, PKM, ENO1) all appear alongside RNA-modifying enzymes, including TRMT61A (Fig. 2N, O). The hypothesis: glycolytic flux regulates tRNA modifications that tune translation. Measuring tRNAs or modifications after glycolytic inhibition would test this metabolic-translational coupling.

These connections represent untested hypotheses generated purely from computational analysis of functional genomics data. While they await experimental validation, they illustrate how DDH can reveal non-obvious metabolic relationships and generate focused, testable predictions that span from nutrient sensing to metabolite transport to cofactor biology.

### Platform Features and User Interface

The datadrivenhypothesis.org platform provides intuitive access to integrated co-essentiality data through a web interface requiring no computational expertise. Users can query individual genes to view dependency distributions across 739 cell lines, identify correlated genes, explore interactive networks, and access pathway enrichments via Enrichr. A co-publication index (0-100 scale) highlights novel relationships—for instance, TP53 and RXFP2 show strong co-essentiality but minimal literature precedent (index = 2), suggesting unexplored biology.

Beyond single genes, users can analyze ∼3,000 predefined pathways from MSigDB or upload custom gene lists. The pathway co-essentiality module generates correlation matrices and hierarchical clustering to reveal functional relationships, as demonstrated by our discovery of 31 pathways uniquely connected to Complex II, including unexpected links to nucleotide and glutamine metabolism.

Quality filters ensure robust results by excluding genes with zero expression (16.8% of gene-cell line pairs) or those present in fewer than 108 cell lines. Statistical thresholds (r > 0.21 or < -0.20) are established through permutation testing. Network topology provides additional validation, where genes appearing in each other’s top correlation lists form high-confidence functional modules, particularly valuable for uncharacterized genes.

The platform offers point-and-click analysis with downloadable results in standard formats, while our GitHub repository (github.com/hirscheylab/ddh) provides complete R pipelines for advanced users. Quarterly updates incorporate new data releases, with version control ensuring reproducibility.

### Integration with the metabolic research ecosystem

DDH complements existing resources while filling critical gaps. Gene expression databases (GEO, ArrayExpress) focus on expression patterns. Protein interaction databases (STRING, BioGRID) capture physical interactions. General co-essentiality browsers (DepMap portal) provide broad analyses. DDH uniquely captures genetic dependencies with pathway-level analysis tailored for metabolic research. The platform integrates with established resources through direct links. HMDB provides metabolite connections. KEGG enables pathway visualization. The Human Protein Atlas shows tissue expression. Specialized portals (T2D Knowledge Portal/dknet.org, Lipid Droplet Database) offer domain-specific information. This integration positions DDH as a central hub for hypothesis generation that connects diverse data types.

### Current limitations and future directions

DDH’s foundation on cancer cell line dependencies represents both a limitation and an opportunity. We acknowledge that transformed cells exhibit altered metabolism: enhanced glycolysis, dysregulated growth signaling, and adapted nutrient utilization patterns, which may not reflect normal physiology. Specialized metabolic functions of differentiated tissues (β-cell insulin secretion, hepatocyte gluconeogenesis, cardiomyocyte fatty acid oxidation) are likely underrepresented or absent. However, this limitation carries an important counterpoint: cancer cells do not invent metabolic pathways *de novo*. Instead, they co-opt, amplify, or rewire existing metabolic networks. The core enzymatic machines, including the electron transport chain, nucleotide biosynthesis, amino acid metabolism, operate through the same fundamental biochemistry whether in a neuron or a lymphoma cell.

Future platform developments are underway to expand beyond the cancer cell paradigm. We are prioritizing integration of data from primary human tissues, including hepatocytes, cardiomyocytes, and immune cells, to complement and contrast with cancer dependencies. This parallel analysis will distinguish fundamental metabolic requirements from cancer-specific vulnerabilities. Additional planned expansions include metabolomics data integration to connect genes with metabolite changes. Via this Forum, we actively seek community contributions of data. These additions will further position DDH as a comprehensive platform for understanding human metabolic gene function across physiological and pathological states.

## Conclusion

datadrivenhypothesis.org transforms the challenge of navigating massive co-essentiality datasets into an opportunity for systematic metabolic gene discovery. By combining pathway-level analysis, multi-dimensional data integration, and intuitive visualization, DDH empowers researchers to generate and test hypotheses about metabolic gene function regardless of their computational expertise. The discovery of the Complex II-purine synthesis axis in AML demonstrates the platform’s potential to reveal clinically relevant metabolic vulnerabilities. As DDH grows through community contributions and quarterly updates, it will continue bridging the gap between big data and biological insight, accelerating our understanding of metabolic networks in health and disease.

## Acknowledgments

We thank the Broad Institute, Human Protein Atlas, Mitocarta, and IEMBase for data, the Duke Center for Genomic and Computational Biology, Dan Leehr, Hilmar Lapp, and OASIS for infrastructure support (JPB), Cédric Scherer for data visualization assistance, and the early DDH users for valuable feedback. This work was supported by NIH grants R01DK115568 and R01AG045351 (MDH), the Duke Cancer Institute P30 Cancer Center Support Grant P30CA014236 (MDH), and a Duke-NUS Block Grant. We also acknowledge Heureka Labs, Inc. for continuing the DDH project. The content is solely the responsibility of the authors and does not necessarily represent the official views of the National Institutes of Health or other funding sources.

## References

1. Li, H., Rukina, D., David, F.P.A., Li, T.Y., Oh, C.M., Gao, A.W., Katsyuba, E., Bou Sleiman, M., Komljenovic, A., Huang, Q., et al. (2019). Identifying gene function and module connections by the integration of multispecies expression compendia. Genome Res 29, 2034–2045. 10.1101/gr.251983.119.

2. Wood, V., Lock, A., Harris, M.A., Rutherford, K., Bahler, J., and Oliver, S.G. (2019). Hidden in plain sight: what remains to be discovered in the eukaryotic proteome? Open Biol 9, 180241. 10.1098/rsob.180241.

3. Yu, M. (2022). Computational analysis on two putative mitochondrial protein-coding genes from the Emydura subglobosa genome: A functional annotation approach. PLoS One 17, e0268031. 10.1371/journal.pone.0268031.

4. Rath, S., Sharma, R., Gupta, R., Ast, T., Chan, C., Durham, T.J., Goodman, R.P., Grabarek, Z., Haas, M.E., Hung, W.H.W., et al. (2021). MitoCarta3.0: an updated mitochondrial proteome now with sub-organelle localization and pathway annotations. Nucleic Acids Res 49, D1541–D1547. 10.1093/nar/gkaa1011.

5. Ferreira, C.R., Rahman, S., Keller, M., Zschocke, J., and Group, I.A. (2021). An international classification of inherited metabolic disorders (ICIMD). J Inherit Metab Dis 44, 164–177. 10.1002/jimd.12348.

6. Wang, T., Yu, H., Hughes, N.W., Liu, B., Kendirli, A., Klein, K., Chen, W.W., Lander, E.S., and Sabatini, D.M. (2017). Gene Essentiality Profiling Reveals Gene Networks and Synthetic Lethal Interactions with Oncogenic Ras. Cell 168, 890–903 e815. 10.1016/j.cell.2017.01.013.

7. Liberzon, A., Subramanian, A., Pinchback, R., Thorvaldsdottir, H., Tamayo, P., and Mesirov, J.P. (2011). Molecular signatures database (MSigDB) 3.0. Bioinformatics 27, 1739–1740. 10.1093/bioinformatics/btr260.

8. Wei, C.H., Allot, A., Leaman, R., and Lu, Z. (2019). PubTator central: automated concept annotation for biomedical full text articles. Nucleic Acids Res 47, W587–W593. 10.1093/nar/gkz389.

9. Stewart, A.E., Zachman, D.K., Castellano-Escuder, P., Kelly, L.M., Zolyomi, B., Aiduk, M.D.I., Delaney, C.D., Lock, I.C., Bosc, C., Bradley, J., et al. (2025). Pathway Coessentiality Mapping Reveals Complex II is Required for <em>de novo</em> Purine Biosynthesis in Acute Myeloid Leukemia. bioRxiv, 2025.2002.2026.640463. 10.1101/2025.02.26.640463.

